# Bacterial communities affected by experimental disturbances to canopy soils of a temperate rainforest

**DOI:** 10.1101/094987

**Authors:** Cody R. Dangerfield, Nalini M. Nadkarni, William J. Brazelton

## Abstract

Trees of temperate rainforests host a large biomass of epiphytes, living plants associated with soils formed in the forest canopy. This study provides the first characterization of bacterial communities in canopy soils enabled by high-depth environmental sequencing of 16S rRNA genes. Canopy soil included many of the same major taxonomic groups of Bacteria that are also found in ground soil, but canopy bacterial communities were lower in diversity and contained different species-level operational taxonomic units. A field experiment was conducted to document changes in the bacterial communities of soils associated with epiphytic material that falls to the forest floor. Bacterial diversity and composition of canopy soil was highly similar, but not identical, to adjacent ground soil two years after transfer to the forest floor, indicating that canopy bacteria are almost, but not completely, replaced by ground soil bacteria. Furthermore, soil associated with epiphytic material on branches that were severed from the host tree and suspended in the canopy contained altered bacterial communities that were distinct from those in canopy material moved to the forest floor. Therefore, connection to the living tree is an important factor in the ecology of canopy soil bacteria. These results represent an initial survey of bacterial diversity of the canopy and provide a foundation upon which future studies can more fully investigate the ecological and evolutionary dynamics of these communities.

**IMPORTANCE:** Temperate rainforests host large accumulations of epiphytes, plants that live on the branches of trees. This study provides a first look at the unique nature of bacterial communities in soil that is formed by the decomposition of epiphytes in the canopy of a temperate rainforest. Falling of epiphytic material results in the transfer of significant amounts of carbon and nutrients from the canopy to the forest floor. This study also experimentally documented how soil bacterial communities are affected by their movement from the canopy to the forest floor. The bacterial composition of canopy soil was altered by the severing of the epiphytic material from the living tree, even when the epiphytes remained in the canopy. Therefore, the unique nature of canopy soil bacteria is determined in part by the living host tree and not only by the physical environmental conditions associated with the canopy.

## INTRODUCTION

Temperate wet forests support a large biomass and high diversity of epiphytes (1–6). These living plants are accompanied by extensive accumulations of organic canopy soils, up to 30 cm thick (3). Single trees can support over 6.5 t dry weight of live and dead epiphytic material (EM), nearly four times the foliar biomass of host trees (5).

These canopy communities play important ecological roles in ecosystem processes, particularly in whole-forest nutrient cycles. Live epiphytic plants derive support from their host trees but acquire nutrients mainly from atmospheric sources (precipitation and particulates that settle within or move through the canopy) (7–9). Canopy soils develop from the accumulation and decomposition of EM on branches and in bifurcations of trees. They consist of the accumulated decomposition products of epiphytic plants, foliage that are derived from mainly allochthonous sources (8–10). Canopy soils retain water and nutrients in their airspaces and on surface exchange sites, respectively (11, 12). When EM falls from branches or “rides down” with broken branches or fallen trees, these nutrients can be transferred to the forest floor and become available to terrestrial vegetation as they die and decompose. Additionally, some host trees gain access to this material directly via canopy roots (11). This material also creates habitat for birds, invertebrates, and arboreal mammals (11, 13–15).

Most studies of EM have focused on their diversity, the pools of nutrients they store, or the ecosystem services they provide. However, little information exists on the biota and processes responsible for the dynamics of EM as it moves from canopy to the forest floor. Epiphytes attached to a fallen tree or branch on the ground may remain vigorous for some time, but the chances for survival of those fallen to the shady ground are low (16). The rates, processes, and biota responsible for their death and decomposition have been documented in only a few tropical forests (17–20). This information is critical to understand the biology and ecological roles of the living communities and their accompanying soils in whole-ecosystem processes.

The difference in environmental conditions between canopy and forest floor has been viewed as the most likely cause of loss of vitality and death of epiphytes when they fall. Different conditions in the soils formed in the canopy vs. forest floor have been documented in a few tropical canopies and temperate rainforests (21–23). In general, canopy soil temperatures are similar to those on the forest floor throughout the year, but canopy soils can experience short, distinct intervals of “dry-downs” during the dry season, which do not occur on the forest floor (23). Other attributes of canopy soils differ from those on the forest floor, e.g., canopy soils are more acidic [canopy pH = 4.6 (3); terrestrial pH=5.4 (24)], and have a higher carbon/nitrogen ratio (3).

These studies and anecdotal observations of epiphyte mortality have lead to the recognition that epiphytes decline and die when they move from the canopy to the forest floor, but the proximate and ultimate factors that contribute to those dynamics are unknown. We carried out an experimental study to explore effects of within-canopy disturbance and movement of EM from the canopy to the forest floor of a temperate rainforest on the resident bacterial communities, which are presumably associated with the decline and decomposition of EM. We compared bacterial community diversity and composition in EM samples that 1) were located on living vs. dead branch substrates; 2) experienced direct vs. indirect contact with the forest floor, with its potentially different cadre of invertebrates, microbes or pathogens; and 3) encountered canopy vs. forest floor environments.

## MATERIALS AND METHODS

### Site Description

The study was conducted in the Upper Quinault River Valley of the Olympic National Park, Washington, USA (47.52°N 123.82°W). Average annual precipitation is ~350 cm in the lowlands and ~510 cm in the higher elevations. The fall, winter, and spring are characterized by heavy rains; summers are typically dry (23). The floodplain forest of the study area is predominated by Big-leaf Maple (*Acer macrophyllum*), which supports the largest epiphyte loads. Other tree species present are Sitka spruce (*Picea sitchensis*), red alder (*Alnus rubra*), and Douglas-fir (*Pseudotsuga menziesii*). Epiphytic material (EM) in Big-leaf maple is described by Aubrey *et al*. (23) and consists of live epiphytes that overlie a thick layer of arboreal soils. Live epiphytes (mosses, liverworts, lichens, and licorice fern, *Polypodium glycyrrhiza)* are dominated by two bryophyte species, (*Isothecium myosuroides* and *Antitrichia curtipendula*). Accumulations of arboreal soils are greatest in branch bifurcations at the trunk (up to 30 cm thick), and taper to small amounts at branch tips. These soils are composed of dead and decomposing epiphytes that remain on host tree branches, and small amounts of intercepted host tree litter. Arboreal and forest floor soil characteristics are described in Tejo et al. (3).

### Sample Collection

On September 28, 2012 we selected nine *A. macrophyllum* trees within previously-established research plots (23). Criteria for inclusion were: safe canopy accessibility; no apparent dead or diseased branches; no visual access from the National Park road; multiple potential sampling branches; mature status; large loads of live epiphytes; and a minimum distance of 200 m from each other. Three of the trees were designated as “experimental trees”, which supported samples of treatments for the experiment. Six were designated as “source trees” in which branches were severed and lowered to the forest floor to provide experimental samples for environmental DNA sequencing studies. From these source trees, 13 branches (6-10 cm in diameter) with complete epiphyte cover were selected, cut, and lowered to the forest floor by an arborist. These branches were cut into 75 cm length segments, and labeled. These severed segments were then randomly selected to one of following treatments within and beneath the experimental trees (**Figure 1**): A) suspended within the canopy (canopy severed, CA-SE), B) placed below on the forest floor (ground perched, GR-PE), or C) EM was stripped and placed directly on the forest floor (ground flat, GR-FL). Two years later (September 14, 2014), we sampled canopy soil from all treatments as well as from undisturbed EM in the canopy of experimental trees (CA-AT) and from ground soil (GR-SO) that was located beneath the drip line of each of the experimental trees (**Figure 1**).

**Figure 1.**
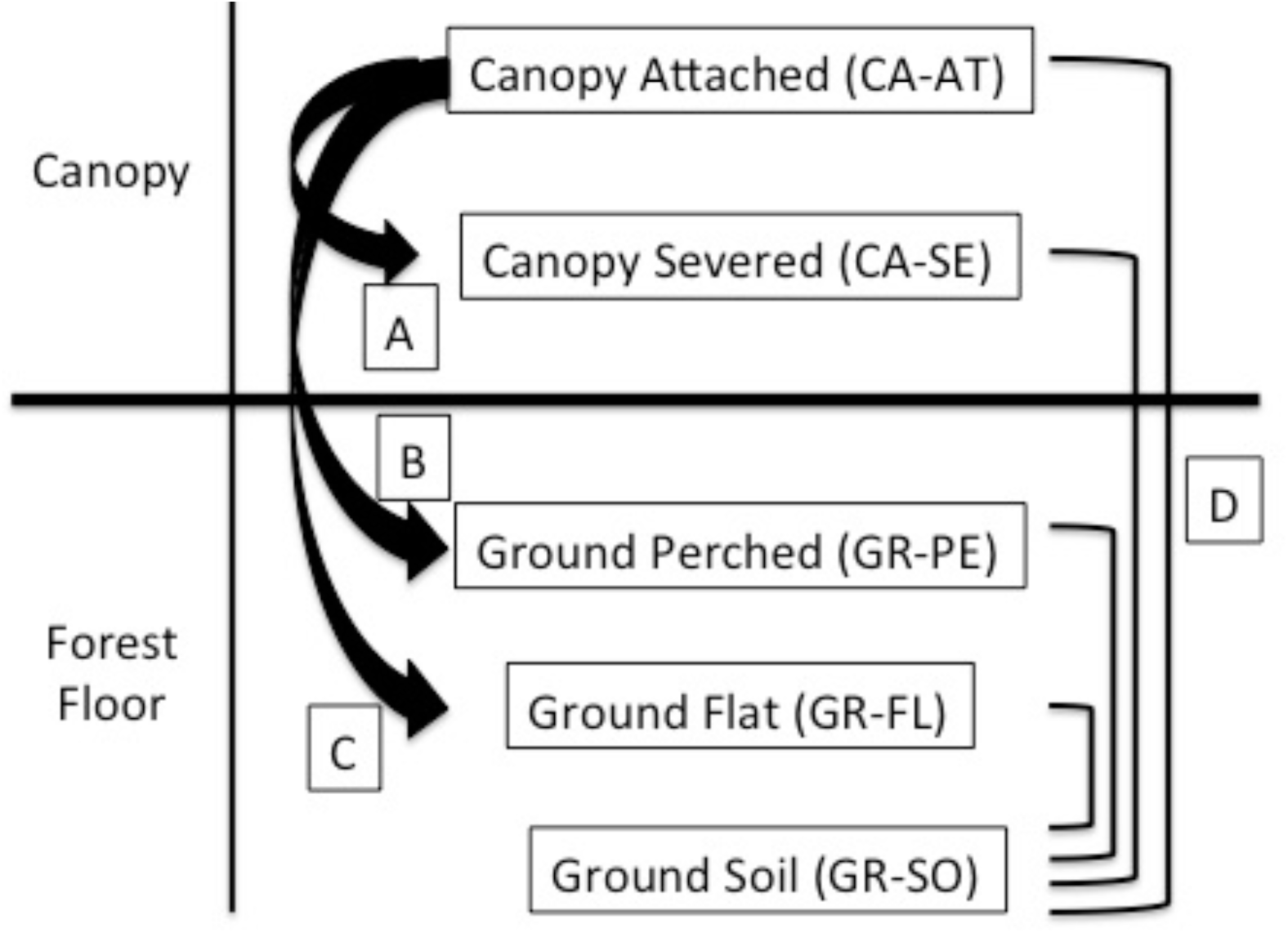
Experimental design to investigate effects of disturbances on soil bacterial community richness, evenness, and composition in the canopy and the forest floor. The undisturbed canopy soil attached to live branches (CA-AT) was compared to three experimental treatments: A) canopy epiphytic material (EM) on severed dead branches suspended in the canopy (CA-SE). B) canopy EM on dead branches transplanted to the forest floor (GR-PE). C) canopy EM removed from the branch and placed directly on the forest floor (GR-FL). In addition, D) all treatments were compared to undisturbed ground soil underneath the tree (GR-SO).

### Extraction of DNA from soil samples

The samples were homogenized for DNA extraction by flash-freezing the sample with liquid nitrogen followed by grinding the sample into a fine powder. DNA was extracted from each sample using the PowerSoil DNA Isolation Kit (MO BIO Laboratories) according to the manufacturer’s instructions and stored at −20°C.

### Bacterial 16S rRNA gene sequencing

Bacterial 16S rRNA gene amplicon sequencing was conducted by the Michigan State University genomics core facility. The V4 region of the 16S rRNA gene (defined by primers 515F/806R) was amplified with dual-indexed Illumina fusion primers as described by Kozich et al. (25). Amplicon concentrations were normalized and pooled using an Invitrogen SequalPrep DNA Normalization Plate. After library QC and quantitation, the pool was loaded on an Illumina MiSeq v2 flow cell and sequenced using a standard 500 cycle reagent kit. Base calling was performed by Illumina Real Time Analysis (RTA) software v1.18.54. Output of RTA was demultiplexed and converted to fastq files using Illumina Bcl2fastq v1.8.4. Paired-end sequences were filtered and merged with USEARCH 8 (26), and additional quality filtering was conducted with the mothur software platform (27) to remove any sequences with ambiguous bases and more than 8 homopolymers. Chimeras were removed with mothur’s implementation of UCHIME (28). The sequences were pre-clustered with the mothur command pre.cluster (diffs=1), which reduced the number of unique sequences from 574,178 to 351,566. This pre-clustering step removes rare sequences most likely created by sequencing errors (29).

### Bacterial Diversity Analyses

These unique, pre-clustered sequences were considered to be the operational taxonomic units (OTUs) of this study and formed the basis of all alpha and beta diversity analyses. We chose not to cluster sequences any more broadly into operational taxonomic units because clustering inevitably results in a loss of biological information and because no arbitrary sequence similarity threshold can be demonstrated to consistently correspond to species-like units. The sample set was reduced from 52 to 37 samples due to extremely low sequence counts in the removed samples suggesting an error had occurred during sequencing. Each of the 37 samples had a minimum of 20,259 sequences, and all samples were randomly sub-sampled down to this minimum prior to calculation of richness, evenness, and alpha diversity. Taxonomic classification of all sequences was performed with mothur using the SILVA reference alignment (SSURefv123) and taxonomy outline (30). Taxonomic counts generated by mothur and edgeR results were visualized in bar charts generated with the aid of the R package phyloseq (31).

### Statistical Analyses

The dissimilarity of bacterial community compositions was calculated with the Morisita-Horn index from a table of OTU abundances across all samples. This index was chosen because it reflects differences in the abundances of shared OTUs without being skewed by unequal numbers of sequences among samples. Morisita-Horn community dissimilarity among samples was visualized with a multi-dimensional scaling (MDS) plot generated with the distance, ordinate, and plot ordination commands in phyloseq. Differences in the relative abundances of sequences between sample types (i.e., categories of samples) were measured with the aid of the R package edgeR (32) as recommended by McMurdie *et al*. (33). The differential abundance of an OTU was considered to be statistically significant if it passed a false discovery rate threshold of 0.05.

### Accession Numbers

All sequence data are publicly available at the NCBI Sequence Read Archive under BioProject PRJNA357844. All SRA metadata, supplementary material, and protocols, are archived with the following DOI: http://doi.org/10.5281/zenodo.208171. All custom software and scripts are available at https://github.com/Brazelton-Lab.

## RESULTS

### Richness and Evenness of Soil Bacterial Communities

**Table 1** lists the operational taxonomic unit (OTU) richness and evenness of bacterial communities inhabiting soil samples collected during this study. EM in the canopy (CA-AT: canopy attached) had lower OTU richness, lower evenness, and lower alpha diversity compared to forest floor soil (GR-SO: ground soil). Bacterial communities of EM on branches that were severed from the tree and suspended in the canopy (CA-SE: canopy severed) had richness and evenness values that were indistinguishable from those of CA-AT samples. Bacterial communities of EM perched on branches that were moved to the forest floor (GR-PE: ground perched) or removed from branches and placed directly on the forest floor (GR-FL: ground flat) had richness and evenness values indistinguishable from those of GR-SO samples. In summary, samples collected from the canopy had lower richness and evenness than samples collected from the forest floor, even if those samples were derived from EM transplanted from the canopy.

**Table 1:** Average Species Richness and Evenness Between Treatments

| Sample Type | S<sub>Obs</sub> | Inverse Simpson | Evenness (from Simpson) |
| --- | --- | --- | --- |
| CA-AT | 9705 ± 2780 | 195 ± 99 | 0.022 ± 0.013 |
| CA-SE | 9464 ± 2349 | 194 ± 143 | 0.022 ± 0.018 |
| GR-PE | 13680 ± 2091 | 678 ± 247 | 0.050 ± 0.021 |
| GR-FL | 13744 ± 2287 | 609 ± 560 | 0.042 ± 0.033 |
| GR-SO | 13168 ± 855 | 561 ± 103 | 0.043 ± 0.010 |

### OTU Composition of Soil Bacterial Communities

At a broad taxonomic level, all samples from canopy and forest floor soils were generally similar, featuring roughly even representation of many bacterial groups commonly found in previously studied soils, including Rhizobiales, Acidobacteria, Actinobacteria, Sphingobacteria, Myxococcales, Xanthomonadales, and Verrucomicrobia. One notable exception is the order Nitrosomonadales (Betaproteobacteria), which was consistently 10-100 times less abundant in canopy compared to ground soils (**Supplementary File S1**). At the level of individual OTUs, differences in bacterial community composition were more easily identified. For example, even though the order Rhizobiales (Alphaproteobacteria) was abundant in both CA-AT and GR-SO samples, the most abundant Rhizobiales OTUs in CA-AT were not abundant in GR-SO (and vice versa). This trend of similar abundances at the phylum, class, and order level but stark contrasts at the OTU level is evident for nearly all of the major taxonomic divisions of Bacteria in the soil samples (**Supplementary File S2**).

In addition to having lower richness and evenness, canopy soils had significantly different OTU compositions compared to ground soils (**Table 2**). The OTU compositions of canopy EM that had been transplanted to the ground (GR-PE and GR-FL), however were very similar to those of undisturbed ground soil (GR-SO) The OTU compositions of CA-SE treatments were highly variable, but their average dissimilarity to CA-AT was greater than the average dissimilarity within CA-SE samples (self-self comparisons in **Table 2**).

**Table 2:** Morisita-Horn Dissimilarity of Bacterial Community Compositions

| DESCRIPTION | COMPARISON | DISSIMILARITY |
| --- | --- | --- |
| Comparison to undisturbed canopy soil | CA-SE vs. CA-AT | 0.765 ± 0.203 |
|  | GR-PE vs. CA-AT | 0.890 ± 0.079 |
|  | GR-FL vs. CA-AT | 0.913 ± 0.094 |
| Comparison to undisturbed ground soil | CA-AT vs. GR-SO | 0.961 ± 0.033 |
|  | CA-SE vs. GR-SO | 0.960 ± 0.023 |
|  | GR-PE vs. GR-SO | 0.704 ± 0.135 |
|  | GR-FL vs. GR-SO | 0.798 ± 0.187 |
| Self-self comparisons | CA-AT vs. CA-AT | 0.628 ± 0.186 |
|  | CA-SE vs. CA-SE | 0.641 ± 0.240 |
|  | GR-PE vs. GR-PE | 0.554 ± 0.136 |
|  | GR-FL vs. GR-FL | 0.955 ± 0.029 |
|  | GR-SO vs. GR-SO | 0.505 ± 0.204 |

Bacterial community dissimilarities were visualized in the MDS plot in **Figure 2**, where each data point represents the OTU structure of one sample and the distance between points represents the dissimilarity between samples. The overall trends evident in **Figure 2** are consistent with the stark contrast between canopy and ground samples shown in **Tables 1-2**. The OTU compositions of most samples collected from GR-PE and GR-FL are more similar to those of GR-SO than to those of CA-AT or CA-SE. **Figure 2** also shows the large variability in the bacterial community compositions of CASE samples and an apparent gradient from the original CA-AT community composition to the most divergent CA-SE community compositions. Furthermore, the shift in CA-SE communities associated with severing the branch from the tree is distinct from the shift in GR-PE and GR-FL communities associated with transplanting the EM from the canopy to the ground (two arrows in **Figure 2**).

**Figure 2.**
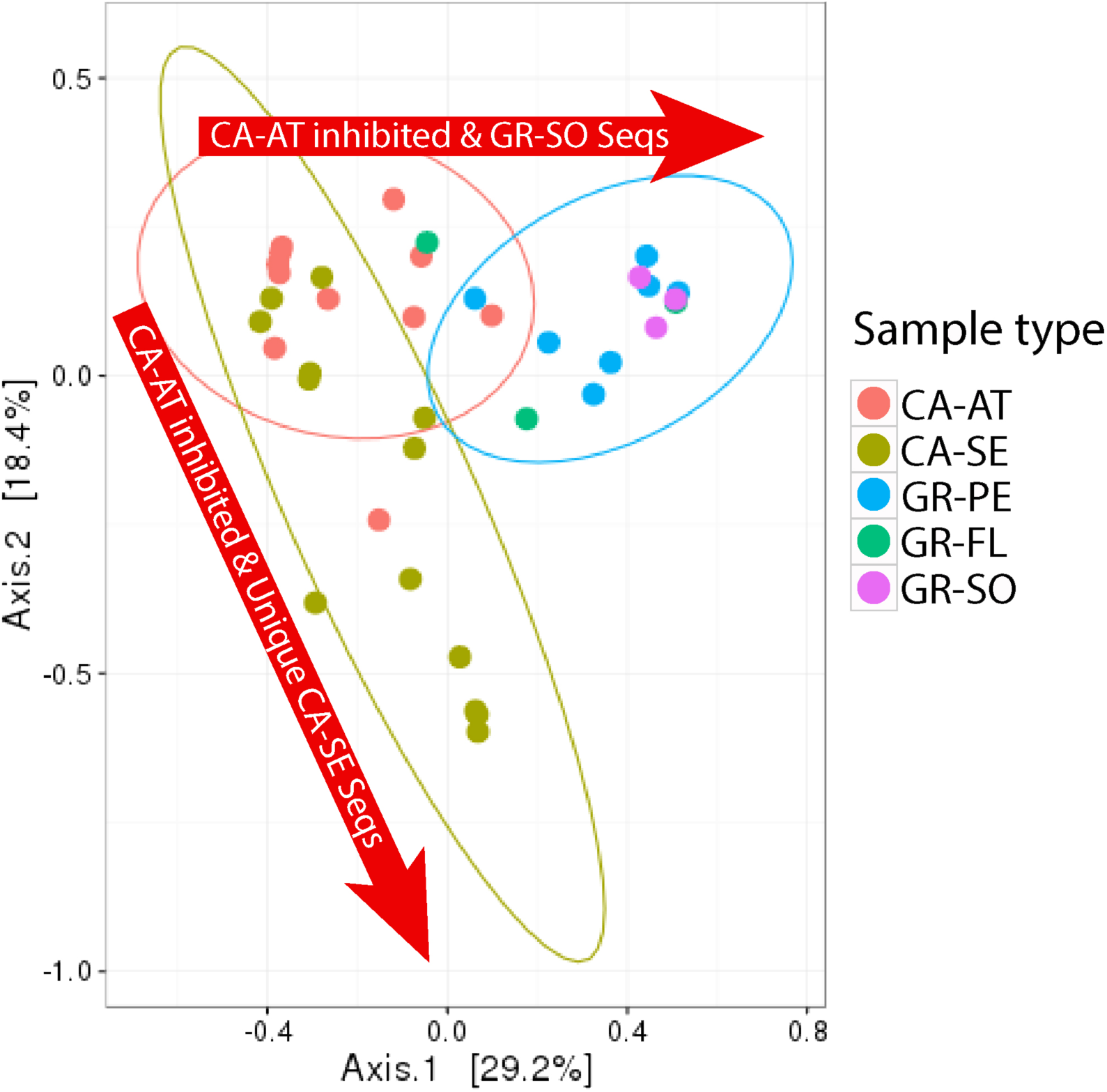
Shifts in bacterial community composition associated with canopy-severed (CA-SE) compared to ground-perched (GR-PE) and ground-flat (GR-FL) treatments with canopy-attached (CA-AT) and ground-soil (GR-SO) representing the original natural community compositions. The ellipses indicate where 95% of samples within a treatment are expected to occur on the plot. Ellipses could only be drawn for sample types containing at least five samples. Arrows reflect the interpretations of which taxa are affected by each treatment, as described in the text.

### OTUs with Differential Abundance in Canopy vs. Ground Soil

To identify individual OTUs that are significantly more abundant in canopy soil compared to ground soil (and vice versa), we contrasted the relative abundances of OTUs in CA-AT to the OTU abundances in GR-SO samples. In **Figure 3**, each data point represents the total abundance of each OTU across all samples (X-axis) and the differential abundance of each OTU between CA-AT and GR-SO (Y-axis). Red data points represent OTUs whose differential abundances passed a significance test (false discovery rate < 0.05) and can be thought of as the OTUs that are characteristic to that sample type. This analysis identified, for example, several Pseudomonadaceae OTUs that were more abundant in ground soil and nearly absent in the canopy (**Figure 3** and **Supplementary File S2**). Furthermore, some OTUs classified as family Bradyrhizobiaceae (order Rhizobiales) were significantly more abundant in ground soil than in canopy soil. The Bradyrhizobiaceae also included other OTUs with the opposite abundance distribution; i.e., they were more abundant in canopy soil than in ground soil. In other words, canopy soil and ground soil each have their own distinct and abundant Bradyrhizobiaceae OTUs. A similar pattern was observed for the Acidobacteriaceae; some OTUs were significantly more abundant in ground soil, and other OTUs were more abundant in the canopy (**Figure 3** and **Supplementary File S2**).

**Figure 3.**
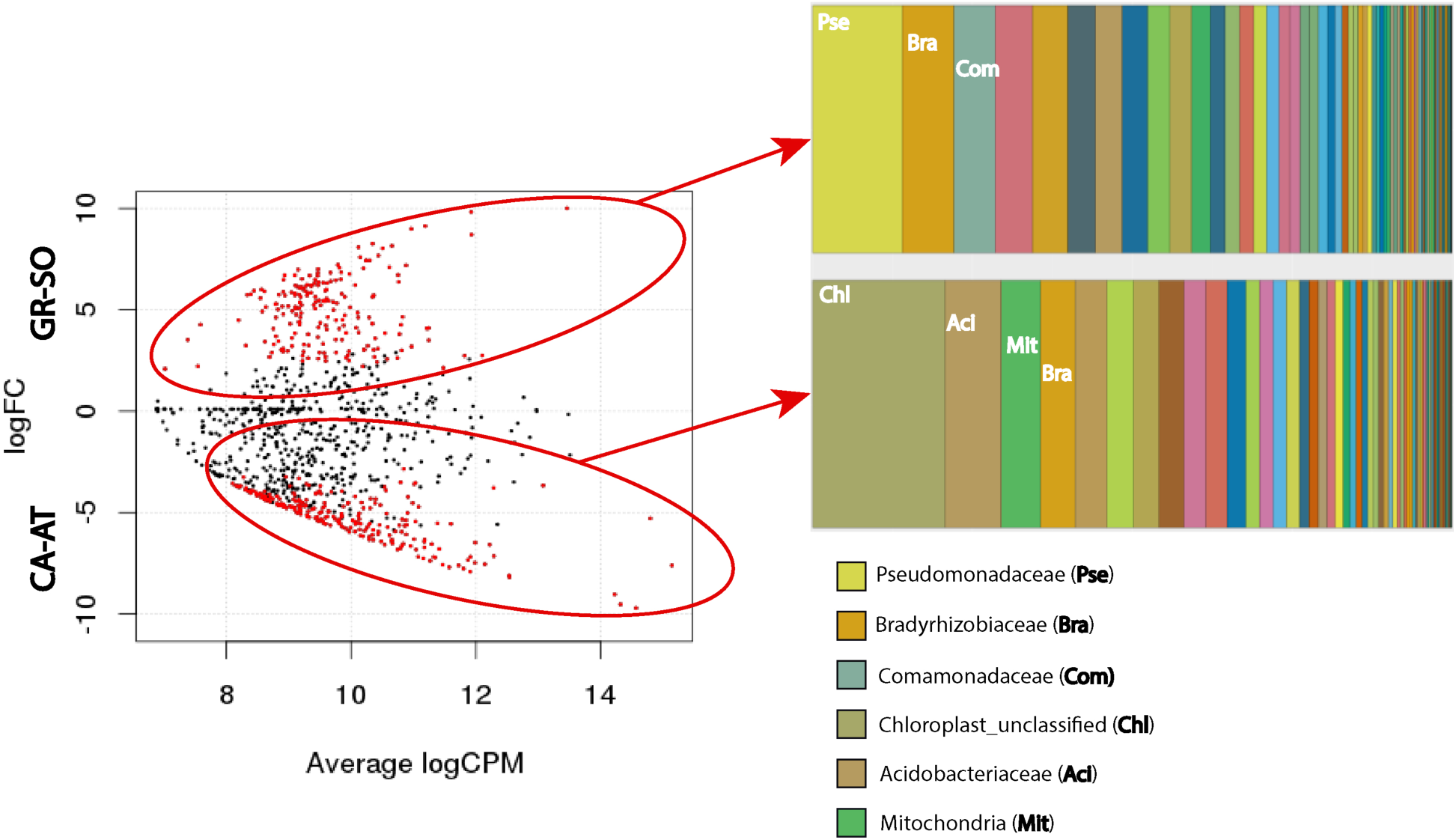
Differential abundance of OTUs in undisturbed canopy soil (CA-AT) and undisturbed ground soil (GR-SO). Red data points indicate OTUs with significantly greater abundance in CA-AT (lower half of plot) or GR-SO (upper half of plot). Significance was defined as false discovery rate < 0.05. Taxonomic classifications of OTUs with differential abundance in each sample type are provided as bar charts. Taxonomic groups with the most numbers of OTUs are labeled with abbreviations defined as bold text in the legend below.

Chloroplasts and mitochondria (both of which are detected by sequencing of bacterial 16S rRNA genes) were among the most common sources of differentially abundant OTUs between CA-AT and GR-SO (**Figure 3**). Most of the chloroplast sequences could not be classified because chloroplast 16S rRNA genes are not reliable taxonomic markers, but the best BLAST hits in the GenBank nonredundant database to the most abundant chloroplast sequences in CA-AT include those belonging to mosses and angiosperms as well as the lycopod *Selaginella*. The most abundant mitochondrial 16S rRNA sequences from CA-AT matched those of diverse ferns, the moss *Funaria hygrometrica*, and the lichenized fungus genus *Psora* (**Figure 3** and **Supplementary File S2**).

### OTUs with Differential Abundance in Experimental Treatments

The abundance distribution pattern of each OTU was examined in order to identify the specific bacterial taxa driving the community shifts associated with experimental disturbances to canopy soil. Nearly all of the highly abundant OTUs were detected in most experimental treatments, but many of these OTUs had significantly greater abundances in one or more treatments compared to CA-AT (red data points in **Figure 4**). There were 164 OTUs more abundant in CA-SE compared to CA-AT (**Figure 4A**), 245 OTUs that were more abundant in GR-PE compared to CA-AT (**Figure 4B**), and 196 OTUs more abundant in GR-FL compared to CA-AT (**Figure 4C**). These differentially abundant OTUs must be primarily responsible for the shifts in community composition evident in **Figure 2**.

**Figure 4.**
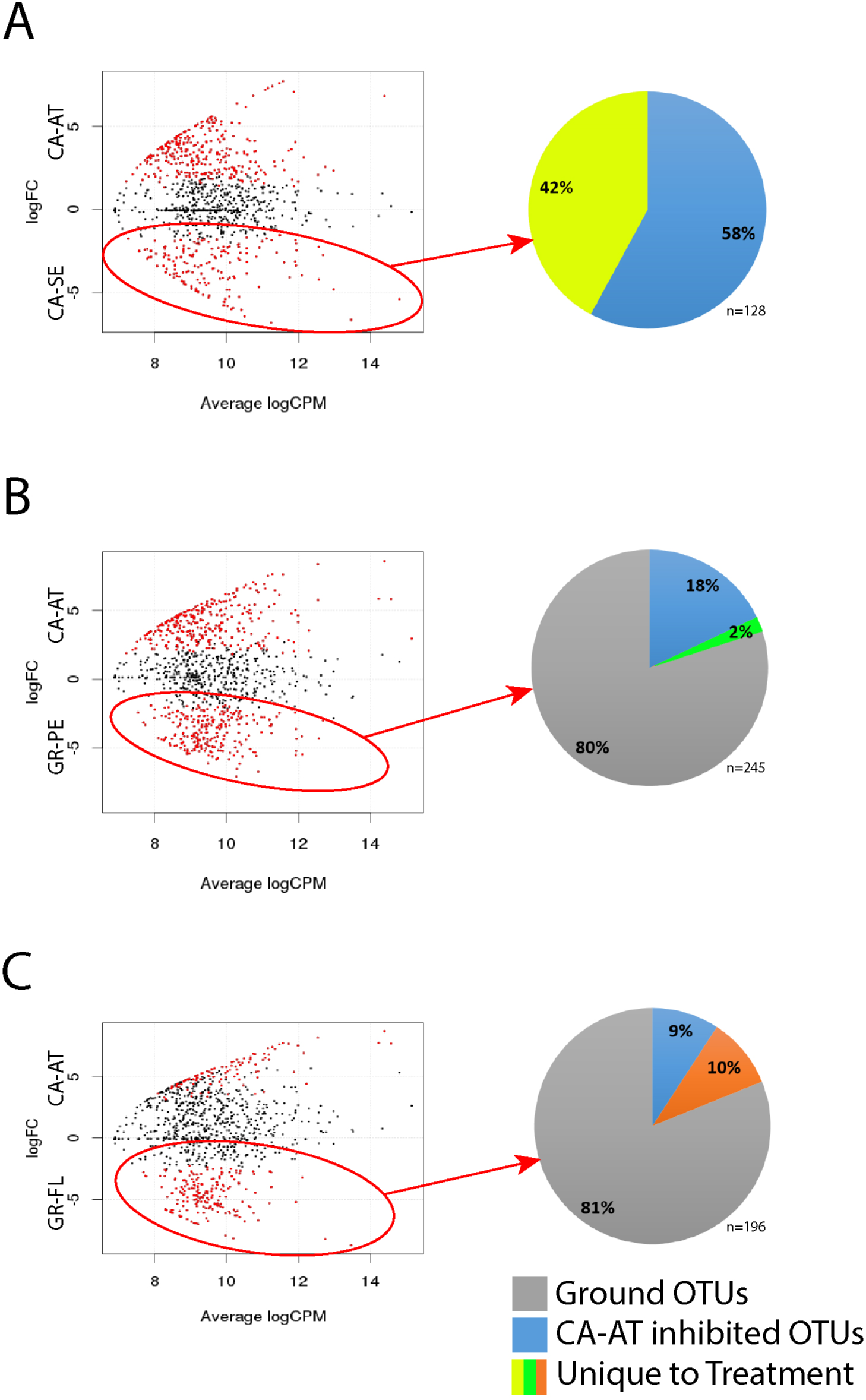
Differential abundance analysis to identify specific taxa with significantly greater abundance in one treatment compared to their abundance in undisturbed canopy soil: (A) CA-SE vs. CA-AT, (B) GR-PE vs. CA-AT, (C) GR-FL vs. CA-AT. Red data points indicate OTUs whose differential abundance passed a significance test (false discovery rate < 0.05). OTUs with significantly greater abundance in disturbance treatments were then categorized by their distribution patterns (shown in pie charts): OTUs that were unique to that treatment, OTUs that were also abundant in nearby ground soil (Ground OTUs), and OTUs that were abundant in all samples except undisturbed canopy soil (CA-AT inhibited).

Most OTUs that were highlighted by the differential abundance tests were found in multiple sample types. For example, 58% of the OTUs that were more abundant in CA-SE compared to CA-AT had similar abundances in GR-PE, GR-FL, and GR-SO (pie chart in **Figure 4A**). Therefore, these OTUs are abundant everywhere except CA-AT and were designated 'CA-AT Inhibited’. The remaining 42% of OTUs that were differentially abundant in CA-SE compared to CA-AT were significantly less abundant or absent in all of the ground samples and were designated ‘Unique CA-SE’.

Nearly all of the OTUs that were differentially abundant in GR-PE and GR-FL compared to CA-AT were found at similar abundances in nearby ground soil (GR-SO). A few of these OTUs were the same OTUs identified as “CA-AT Inhibited” above, and the remaining OTUs were designated as “Ground OTUs” (pie charts in **Figures 4B-4C**), which are inferred to be derived from the nearby ground soil. Very few OTUs were uniquely abundant in the GR-PE or GR-FL treatments, which is consistent with the positions of GR-PE and GR-FL samples overlapping with those of CA-AT and GR-SO samples in the MDS plot of **Figure 2**.

### Taxonomic Classifications of Differentially Abundant OTUs

In general, the differentially abundant OTUs included representatives from all of the typical soil taxonomic groups listed above and were not obviously divergent from the general population at broad taxonomic levels. A notable exception is that OTUs classified as family Acidobacteriaceae (phylum Acidobacteria) and family Acidothermaceae (phylum Actinobacteria) were much more abundant in CA-AT compared to any of the treatments (**Supplementary File S2**).

The “CA-AT Inhibited” and “Unique CA-SE” categories of OTUs were also similar at broad taxonomic levels but differed at more specific taxonomic resolution (**Supplementary File S2**). For example, all Rhizobiales OTUs that were more abundant in CA-SE than CA-AT and classified as family Bradyrhizobiaceae (including genus *Bradyrhizobium*, which is typically found in plant root nodules) were identified as “CA-AT Inhibited” because these sequences were also abundant in ground soil (GR-SO). In contrast, several unclassified Rhizobiales OTUs in CA-SE were absent in ground soil and were therefore included in the “Unique CA-SE” category. Within phylum Actinobacteria, OTUs in class Actinobacteria were overwhelmingly “CA-AT Inhibited” while class Thermoleophilia were mostly “Unique CA-SE”. OTUs classified as Xanthomonadales were also found in both “CA-AT Inhibited” and “Unique CA-SE” categories. Chloroplast and mitochondria sequences with high abundance in CA-SE were mostly absent in GR-SO (and are therefore included in the “Unique CA-SE” category), and many of these sequences were similar to those from mosses and liverworts (**Supplementary File S2**). The “CA-AT Inhibited” category also included many Chloroflexi OTUs (classes Anaerolineae and Ktedonobacteria).

## DISCUSSION

### The Unique Bacterial Communities of Canopy Soil

Canopy soils are assumed to be a harsh environment for most microorganisms, due to their higher acidity (3) and to the periodic “dry-downs” during the summer (23). Our results demonstrate that the bacterial communities of canopy soils have much lower diversity than those in ground soils (**Table 1**). Nevertheless, this lack of diversity is not reflected in a dramatically different community composition. Rather, the bacterial community compositions of canopy soils are recognizably similar to those of ground soils. All of the major taxonomic groups of Bacteria found in the soil of the forest floor were also identified in canopy soils.

Cataloguing individual OTUs that responded to experimental disturbances of EM provided deeper insights into the distinct nature of canopy soils. The most abundant ‘missing microbes’ of the canopy (i.e., those contributing to the lower diversity in the canopy) were identified as a set of “CA-AT Inhibited” taxa that were prevalent in all experimental treatments but not in the original, undisturbed canopy soil. The taxonomic classifications of the “CA-AT Inhibited” taxa are not clearly distinct from the general population. For example, some of the most abundant OTUs belong to the Actinobacteria and Bradyrhizobiaceae, which are also represented in the canopy, but by different and many fewer OTUs. These results are consistent with the “CA-AT Inhibited” taxa representing widespread soil bacteria that are unable to thrive in the harsh conditions of the canopy.

### Ground Soil Bacteria Dominate Canopy Material Transplanted to the Forest Floor

The deposition of canopy EM onto the forest floor must stimulate microbial activity, which could be associated with colonization of the EM by nearby ground soil organisms, or by stimulation of organisms that are already present in the canopy EM, or both. Although disentangling cause and effect is not possible with the available data, our results yield insights into the dynamics of bacterial populations in response to disturbances of the canopy EM. First, degradation of canopy EM on the forest floor is accompanied by a replacement of canopy bacteria with typical ground soil bacteria such that the community composition is highly similar to nearby ground soil within two years (**Figure 2**). Second, this transition to a typical ground soil community appears to be unaffected by whether the canopy material is retained on or removed from the branch. Third, our results provide very little evidence that the movement of EM to the forest floor stimulated the growth of bacteria that were native to the canopy. Such organisms would have been detected as OTUs with greater abundance in the transplanted material compared to the original canopy soil and also compared to the ground soil. Very few such OTUs were identified (labeled “Unique to Treatment” in **Figure 4**). In contrast, the vast majority of OTUs in the transplanted EM could be traced to nearby ground soil (“Ground OTUs” in **Figure 4**).

These results suggest that the accelerated degradation of canopy soils when placed on the forest floor is caused primarily by colonization of the canopy material by nearby ground soil bacteria. However, stimulation of resident bacteria could also play a role, considering that the transplanted materials still retained a few unique taxa that could not be traced to ground soil, suggesting that the legacy of the canopy is still evident in these samples. Additional work is needed to test whether this is a consistent signal or simply due to incomplete sampling.

### Severing the Connection to the Living Tree Causes Distinct Shifts in the Bacterial Community

Canopy soils on branches that were severed from the host tree and suspended in the canopy for two years contained bacterial communities that were distinct from the original canopy community and also from ground soil. These distinctive bacterial communities could have arisen due to dispersal of bacteria from ground soil or from another source not captured by the experimental design. These possibilities were evaluated by categorizing all bacterial OTUs with greater abundance in the severed branch into two groups: those that are also abundant in ground soil (“CA-AT Inhibited”) and those not detected at high abundance in any other samples (“Unique CA-SE”). These two groups contained roughly equal numbers of abundant OTUs, indicating that the bacteria able to thrive in the severed branch are derived from ground soil as well as other undetermined areas of the forest.

Canopy material in the severed branch did not experience accelerated degradation, unlike the material transplanted to the forest floor. However, during visits to the canopy during the study period, EM on severed branches appeared to be drier than EM on intact branches, perhaps because the severed branches could not receive stemflow. These observations, together with the bacterial diversity results, suggest that the severed branches are even harsher environments than intact branches of the canopy where some, but not all, of the original canopy community persists and is altered by the colonization of a few opportunistic taxa from elsewhere in the forest.

### Conclusions

Epiphytic material and associated soils in the canopy constitute large pools of nutrients, water, and carbon in temperate rainforests (3, 34). Therefore, the origin and fate of canopy epiphytic material is of central importance to understanding the microbial ecology of temperate rainforests. Our results provide the first in-depth survey of bacterial communities in canopy soils and reveal them to be taxonomically similar to underlying ground soil but much lower in diversity. The comparatively few bacterial taxa that are highly abundant in canopy soil are distinct members of the same taxonomic groups found in ground soil. Our field experiment demonstrated that the soil created by EM decomposing on the forest floor for two years is nearly indistinguishable from ground soil. However, epiphytic material in the canopy that has been severed from the host tree fosters unique and low-diversity bacterial communities. Many of the bacterial taxa stimulated in the severed branch are widely distributed among the canopy and forest floor, suggesting that they might be exploiting an opportunity to colonize a habitat that has just experienced a massive disturbance. These results highlight the unique nature of canopy-dwelling bacterial communities as well as the importance of the connection to a living tree as an essential component of their canopy ecology.

## Acknowledgements

We thank Alex Hyer, Christopher Thornton, Emily Dart, and August Longino for technical assistance with laboratory work and data analyses. Erika Longino, Dennis Aubrey, Autumn Amici, Camila Tejo, Jordan Herman, and Johanna Castillo provided help in the field. Jerry Freilich provided research administrative support in the Olympic National Park. The research was partially funded by grants from the National Science Foundation (DRL 15-14494 and DEB 11-41833) and the University of Utah. Research permits were issued from the Research Office at the Olympic National Park (OLYM-000234).

## Supplementary Material

**Supplementary File S1**: Taxonomic classifications of bacterial OTUs provided as an html file that displays taxonomy as an interactive krona plot for each of the 37 samples.

**Supplementary File S2**: Abundance, taxonomic classification, and distribution category of each OTU represented by ≥10 sequences across all samples.

